# Exploring Transcriptomic Changes in Antibiotic-Resistant *Salmonella* Heidelberg Modulated by Commensal *Escherichia coli* in Vitro

**DOI:** 10.1101/2024.12.12.628196

**Authors:** Yasir R. Khan, Lekshmi K. Edison, Thomas Denagamage, Subhashinie Kariyawasam

## Abstract

Nontyphoidal *Salmonella* (NTS) are major foodborne pathogens primarily transmitted to humans through contaminated poultry products. The increase in antibiotic-resistant NTS, including *Salmonella* Heidelberg, has recently become a public health issue. Current control measures are inadequate, emphasizing the need for novel approaches to mitigate NTS colonization in poultry and contamination of poultry products. We hypothesized that commensal *Escherichia coli* can reduce antibiotic-resistant NTS colonization in the chicken intestines by modulating the fitness, virulence, and antibiotic resistance (AMR) potential of *Salmonella*. To test this hypothesis, we co-cultured commensal *E. coli* 47-1826 and antibiotic-resistant *S*. Heidelberg 18-9079 strains isolated from poultry and analyzed their transcriptomes using RNA sequencing. Our analysis revealed 4,890 differentially expressed genes in *S*. Heidelberg 18-9079 when co-cultured with *E. coli* 47-1826. After filtering the expression data, we found 193 genes were significantly upregulated while 202 genes were downregulated. Notably, several genes involved in bacterial growth, pathogenicity and virulence, biofilm formation, metal-ion hemostasis, signal transduction and chemotaxis, stress response, transmembrane transport of xenobiotics, and cellular metabolism were down-regulated by as much as eighty-six folds in *S*. Heidelberg as compared to the control. Further, the study revealed the downregulation of genes associated with AMR and drug efflux in *S.* Heidelberg 18-9079 by up to twelvefold. These findings highlight that commensal *E.* c*oli* 47-1826 may reduce the fitness, persistence, virulence, and antimicrobial resistance (AMR) dissemination of *S.* Heidelberg 18-9079, implying that *E. coli* strain 47-1826 could be utilized as a strategy to mitigate antibiotic-resistant S. Heidelberg in poultry, ultimately enhancing food safety.

**Importance:** Nontyphoidal *Salmonella*, commonly transmitted to humans through contaminated poultry meat and eggs, is one of the most frequent causes of foodborne illness. Augmenting the situation, foodborne outbreaks of antibiotic-resistant NTS have become an additional food safety and public health concern. Evaluation of growth changes and transcriptomic profiling of antibiotic-resistant *S.* Heidelberg and commensal *E. coli* in a mixed culture of the two will provide insight into the ability of commensal *E. coli* to reduce *S*. Heidelberg colonization of chicken intestines and the genes involved in that change. Our study showed that commensal *E. coli* strain 47-1826 significantly reduced antibiotic-resistant *S.* Heidelberg 18-9079 counts and expression of *Salmonella* genes, which play a vital role in the growth and persistence of nontyphoidal *Salmonella* in poultry intestines. The study results suggest the potential use of commensal *E. coli* strain 47-1826 to control antibiotic-resistant *S.* Heidelberg colonization in poultry, leading to improved food safety through reduced NTS contamination of foods of poultry origin and reduced dissemination of antibiotic-resistant *Salmonella* and their resistant determinants to humans via the food chain.

## 1. Introduction

Nontyphoidal *Salmonella* (NTS) are a frequent cause of bacterial foodborne illness in humans globally (CDC 2024). Among the NTS implicated in foodborne illness, *Salmonella enterica* serovar Heidelberg (*S*. Heidelberg) has been documented as the serovar associated with the second-highest *Salmonella-*related mortalities, more widespread foodborne outbreaks, and resistance to antibiotics critical to human health (Hoffmann et al., 2012; Denagamage et al., 2019). The National Antimicrobial Resistance Monitoring System (NARMS) has indicated multidrug resistance (MDR) approximately in one-fifth of human isolates of *S*. Heidelberg in 2021 in the United States (CDC, 2024). Similarly, a significant increase in MDR in *S*. Heidelberg isolates over the last two decades is also a serious concern. *S*. Heidelberg is frequently isolated from food products of animal origin, and poultry is considered as one of the primary reservoirs of antibiotic-resistant *Salmonella* and their resistance gene repertoire (Hoffmann et al., 2012). The more invasive and severe forms of *S*. Heidelberg infections in humans compared to other *Salmonella* serovars project its presence in poultry production systems as a critical problem due to the increased risk of contamination of poultry commodities, and subsequent transmission to humans (Taylor et al., 2015). Furthermore, owing to the ease of dissemination and continuous increase in the virulence, the spread of *S*. Heidelberg mainly through poultry products is considered a significant public health and food safety issue (Nair et al., 2021).

Many countries, particularly the developing countries, still use antibiotics to control *Salmonella* in poultry, raising public health concerns and contributing to antibiotic resistance (AMR) in *Salmonella* serovars (Fagbamila et al., 2023). The emergence of MDR in *S.* Heidelberg, especially against extended-spectrum cephalosporins (ESCs) has largely reduced the treatment options for invasive *S*. Heidelberg infections in humans (Han et al., 2012; Hoffman et al., 2012). However, inconsistent outcomes and limited effectiveness of various pre-harvest and post-harvest approaches to minimize *Salmonella* colonization and persistence in the poultry intestines, their dissemination in the environment, and contamination of the food chain emphasize the need for more effective and reliable interventions across the poultry production continuum (Sun et al., 2022; Dewi et al., 2021; Nair et al., 2021). Among many approaches investigated, developing non-toxic, residue-free, potent agents to control *Salmonella* colonization in poultry has been the focus of contemporary efforts. Such microbial agents, called probiotics, are now under consideration because of their effectiveness against pathogen colonization in the intestines, and additional beneficial effects on the host (Chen et al., 2020).

Probiotics have been shown to enhance resistance to *Salmonella* colonization in the poultry gut (Dewi et al., 2021). In this context, commensal *E. coli* can be the key bacterial species to effectively control the intestinal colonization of *Salmonella* in chickens (Litvak et al., 2019). Commensal *E. coli* living symbiotically in the chicken gut are involved in several functions, including intestinal cell maturation and development, enzyme production, nutrients synthesis, immunomodulation, homeostasis, competitive expulsion of pathogens, and modulation of the intestinal microbiome (Patterson & Burkholder 2003). Because intestinal commensal *E. coli* co-exists with *Salmonella* in the intestinal tract of chickens, efforts are underway to identify specific strains of commensal *E. coli* as probiotics against NTS serovars in poultry (Deriu et al., 2013; Sun et al., 2022; Wu et al., 2023). However, the effect of intestinal commensal *E. coli* of poultry origin on the colonization, persistence, virulence, and AMR dissemination of foodborne antibiotic-resistant *S*. Heidelberg in the chicken intestinal tract has not yet been fully elucidated.

Transcriptomic profiling provides global gene expression patterns central to understanding the complex interplay between microbial populations. Comparative transcriptomic analysis further identifies differentially expressed genes and pathways under specific conditions, providing a holistic insight into the molecular basis for interactions between the host and pathogen or among the members of different microbial communities (Edison et al., 2024). However, literature describing the comparative transcriptomic profiling of *S*. Heidelberg in the presence of commensal *E. coli* remains scarce. Therefore, to narrow this knowledge gap, the current study examined the transcriptomic profile of a strain of *S*. Heidelberg in the presence of an intestinal commensal strain of *E. coli.* Here, we hypothesized that intestinal commensal *E. coli* 47-1826 of poultry origin could reduce the expression of genes critical for fitness, virulence, and dissemination of AMR in antibiotic-resistant *S*. Heidelberg18-9079 strain. Through whole genome transcriptomic profiling, we investigated the differential gene expression of *S.* Heidelberg 18-9079 in the presence of commensal *E. coli* 47-1826 and explored the effects of commensal *E. coli* 47-1826 on the fitness, virulence, and AMR dissemination potential of *S*. Heidelberg 18-9079 to use it as a pre-harvest intervention strategy to mitigate intestinal colonization of NTS in poultry.

## 2. Results

### 2.1. Transcriptomic profile of *S.* Heidelberg 18-9079 and commensal *Escherichia coli* 47-1826 in co-culture

The transcript reads were mapped to *Salmonella* enterica subsp. *enterica* serovar Heidelberg (GCF_016452005.1) and ESC-resistance encoding plasmid (NZ_MW 349106.1) for *S*. Heidelberg 18-9079, and *E. coli* APEC O1 (GCF_000014845.1) for commensal *E. coli* 47-1826. The percentage map read coverages of transcript reads were 99.64 and 99.31 for *S*. Heidelberg 18-9079 and *E. coli* 47-1826 grown individually (control), while 99.47 and 99.52 for *S*. Heidelberg 18-9079 and *E. coli* 47-1826 in the co-culture, respectively. Diagnostic plots for the read count data were generated to analyze the variations in mapped library sizes of significantly differentially expressed genes (SDEGs) for *S.* Heidelberg 18-9079 (Figure 2a, 2c, 2e) and commensal *Escherichia coli* 47-1826, respectively (Figure 2b, 2d, 2f).

**Figure 1:**
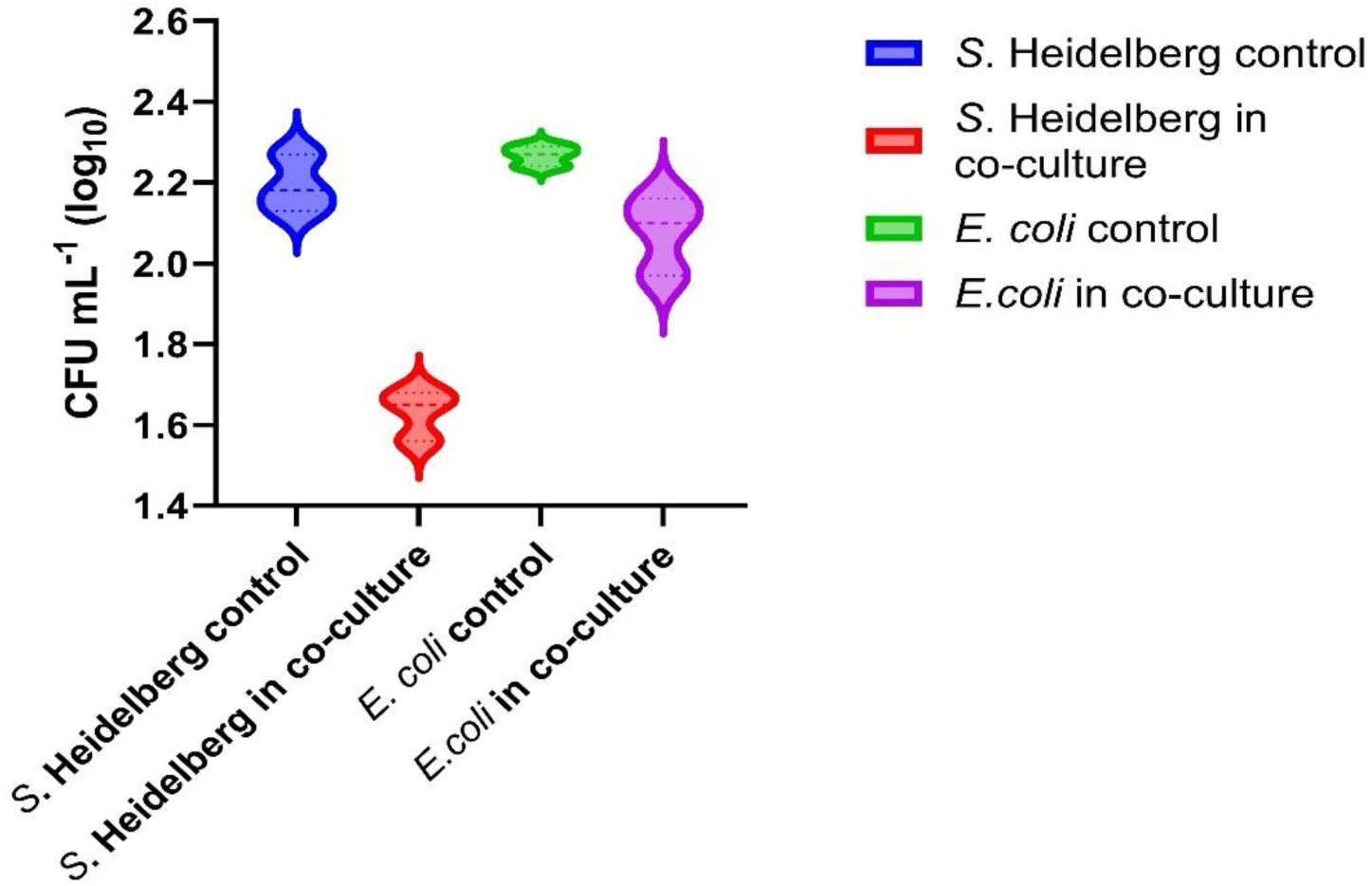
Colony forming units (mean ± SD × log_10_ CFU/mL) of *S*. Heidelberg and commensal *E. coli* in co-culture and controls. The log_10_ CFU/mL of *S*. Heidelberg in the co-culture was significantly less than the control (p ≤0.05).

**Figure 2:**
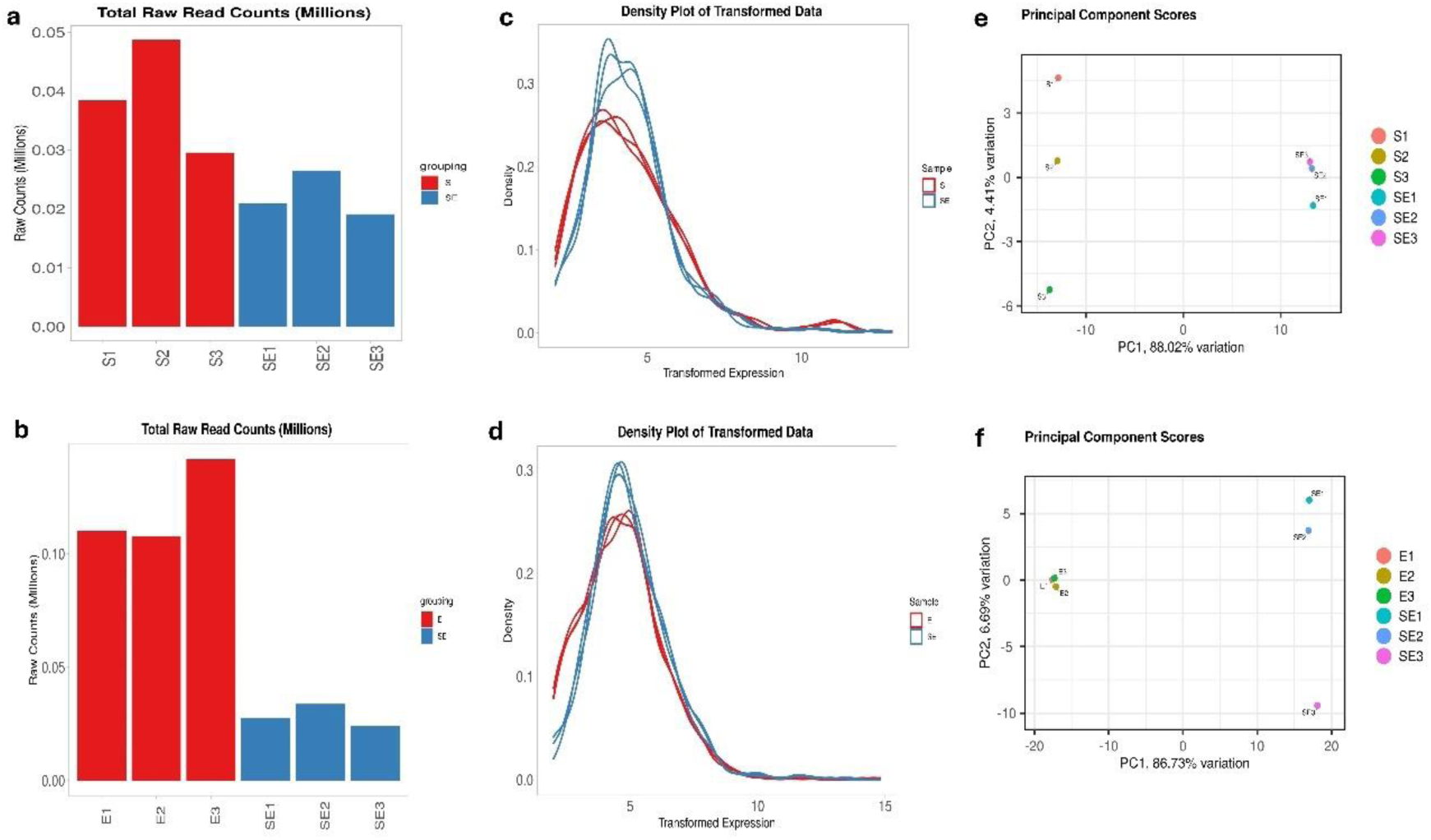
Diagnostic plots illustrating the sequencing depth bias for *S*. Heidelberg 189079 and commensal *E. coli* 47-1826 grown alone (controls) and co-culture. Total raw read count per library (a, b), density plot of regularized log-transformed total read counts (c, d) and principal component analysis (PCA) (e, f) of *S*. Heidelberg 18-9079 and commensal *E. coli* 47-1826, respectively. The difference between the *S.* Heidelberg 18-9079 (S1, S2, S3) grown alone (control) and *S*. Heidelberg 18-9079 in co-culture with *E. coli* 47-1826 (SE1, SE2, SE3) was accounted for the observed 88.02% variation in *S.* Heidelberg 18-9079 in the co-culture compared to the control (Figure 2e). The total read counts differed significantly among the sample groups (p= 4.91e-02), based on ANOVA. Total read counts differed significantly different among sample groups (p= 1.26e-03) based on ANOVA.

### 2.2. Overall transcriptomic differences indicated downregulation of *S.* Heidelberg 18-9079 genes responsible for propagation, persistence, virulence, and antibiotic resistance dissemination

Transcriptomics profile of *S*. Heidelberg 18-9079 co-cultured with *E. coli* 47-1826 differed from *S*. Heidelberg 18-9079 grown alone, and gene expression analysis showed a significant downregulation of genes critical for *S*. Heidelberg 18-9079 propagation, persistence, virulence, and AMR dissemination in the presence of commensal *E. coli* 47-1826. Results of the overall differential gene expression analysis of *S*. Heidelberg 18-9079 when co-cultured with *E. coli* 47-1826 (*S*. Heidelberg 18-9079 *+ E. coli* 47-1826 vs. *S*. Heidelberg 18-9079) are presented in supplementary file S2. Briefly, among 4,968 differentially expressed genes (DEGs) in *S*. Heidelberg 18-9079 when co-cultured with *E. coli* 47-1826, 2158 genes were downregulated, and 2230 were upregulated, whereas 580 hypothetical genes exhibited no detectable change. However, after applying gene filtration criteria, only 395 of 4968 genes remained as significantly differentially expressed genes (SDEGs) in *S*. Heidelberg 18-9079. Of these 395 SDEGs of *S*. Heidelberg 18-9079, 193 genes were downregulated, and 202 upregulated. Overall differential transcriptomic changes in *S*. Heidelberg 18-9079 are presented in volcano plot (Figure 3a) and heatmaps of the expression profiles of different SDEGs categories are presented in (Figure 4).

**Figure 3:**
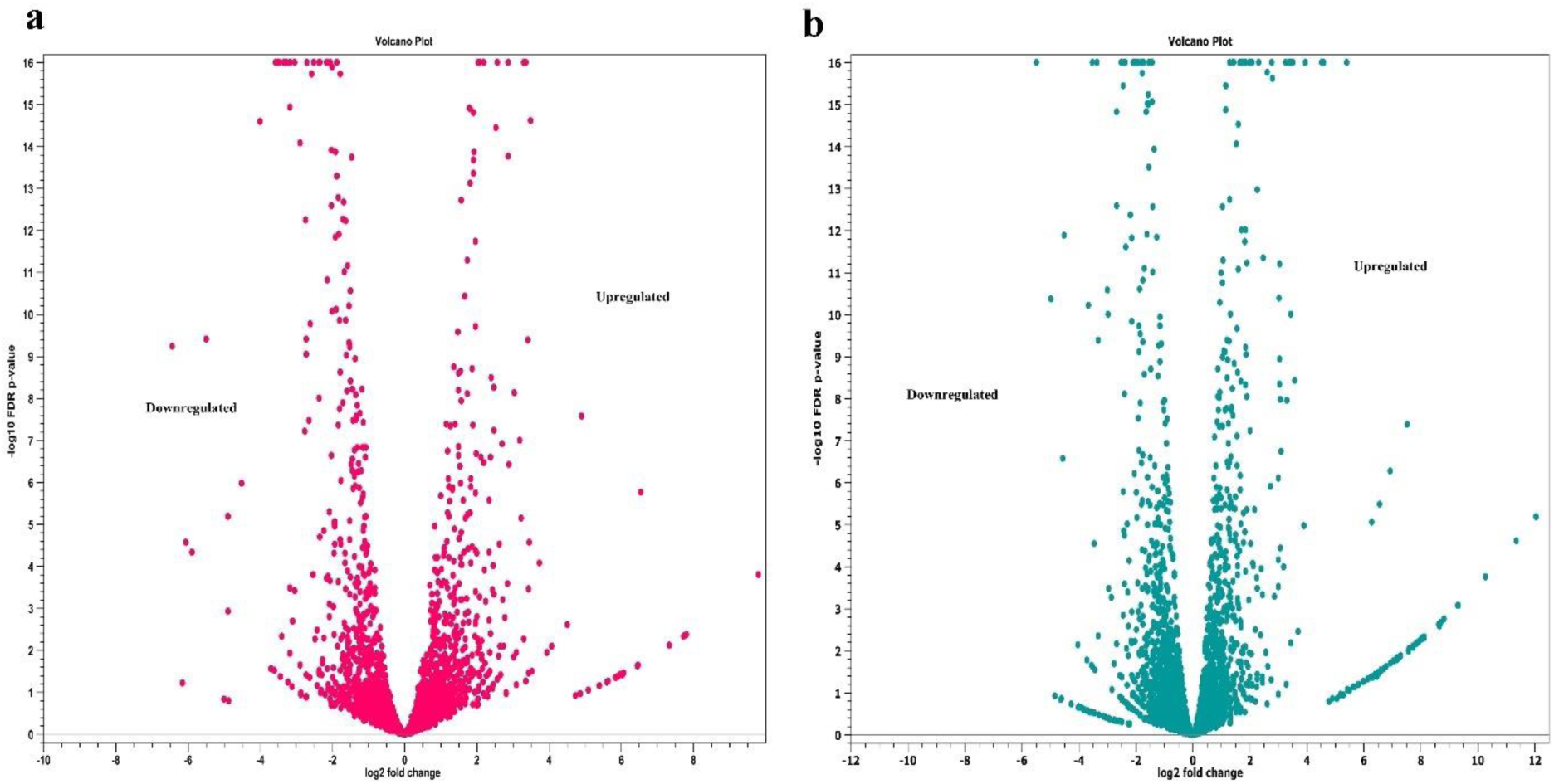
Volcano plots depicting the differential transcriptomic profile of *S.* Heidelberg 18-9079 (a) and commensal *E. coli* 47-1826 (b) in co-culture. This figure provides the relationship between the magnitude (X axis = log_2_ fold change) and extent of gene expression (Y axis = p-value) in the overall transcriptomic expression of *S.* Heidelberg 18-9079 and commensal *E. coli* 47-1826, respectively.

**Figure 4:**
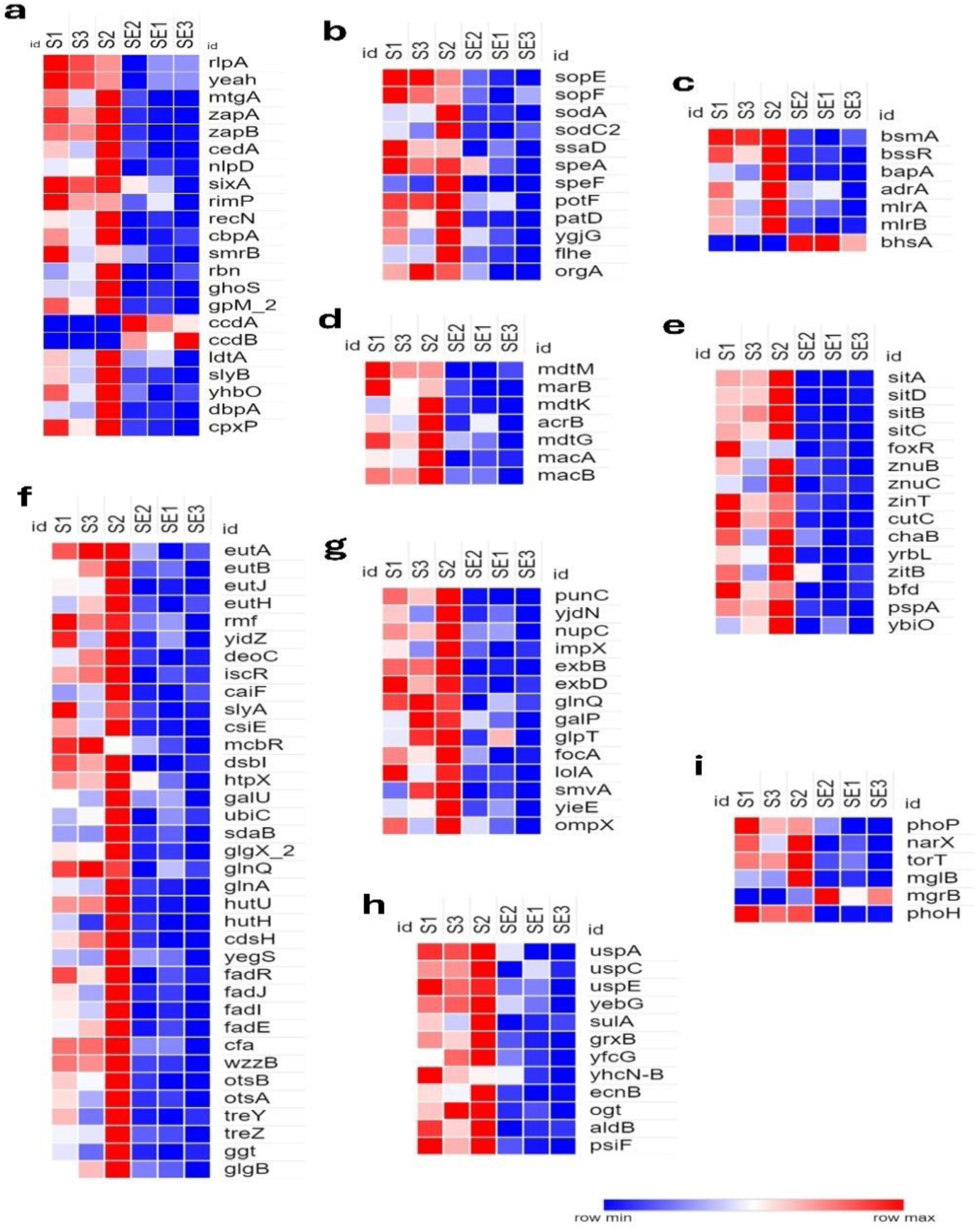
Heat maps of transcriptional responses of *S.* Heidelberg18-9079 when co-cultured with *E. coli* 47-1826 based on functional analysis. (**a**) cell proliferation, (**b**) pathogenicity and virulence factors, (**c**) biofilm formation, (**d**) antimicrobial resistance and multidrug efflux, (**e**) ion homeostasis, (**f**) cellular metabolism, (**g**) outer membrane proteins and transmembrane transporters, (**h**) stress response regulation, and (**i**) signal transduction and chemotaxis. The transformed Fragments/Kilobase of Transcript/Million (FPKM) numbers (normalized read counts) are illustrated in color. Blue indicates low expression, whereas red indicates high expression.

### 2.3. Overall transcriptomic differences showed upregulation of *E. coli* 47-1826 genes responsible for propagation, adherence and motility, metal ion homeostasis, stress response regulation, and signal transduction

A significant upregulation of genes important for propagation, adherence and bacterial motility, ions homeostasis, signal transduction and chemotaxis, stress management, transmembrane transportation, and cellular metabolism, was observed in *E. coli* 47-1826 on gene expression analysis in the presence of antibiotic-resistant *S*. Heidelberg 18-9079. Briefly, among 5227 DEGs in *E. coli* 47-1826 in co-culture with *S*. Heidelberg 18-9079, 2023 genes were upregulated, 2021 genes downregulated, and 1183 hypothetical genes exhibited no detectable change. However, 660 genes were SDEGs out of 5227 genes in *E. coli* 47-1826 after the implementation of gene filtration criteria. Of these 660 SDEGs, 237 genes were upregulated, and 423 were downregulated. Overall differential transcriptomic changes in *E. coli* 47-1826 are presented in volcano plot (Figure 3b). Heatmaps of the expression profiles of different SDEGs categories are presented in (Figure 5). Results of the overall differential gene expression analysis of *E. coli* 47-1826 when co-cultured with *S*. Heidelberg (*E. coli* 47-1826 vs. *S*. Heidelberg 18-9079 *+ E. coli* 47-1826) are presented in S3.

**Figure 5:**
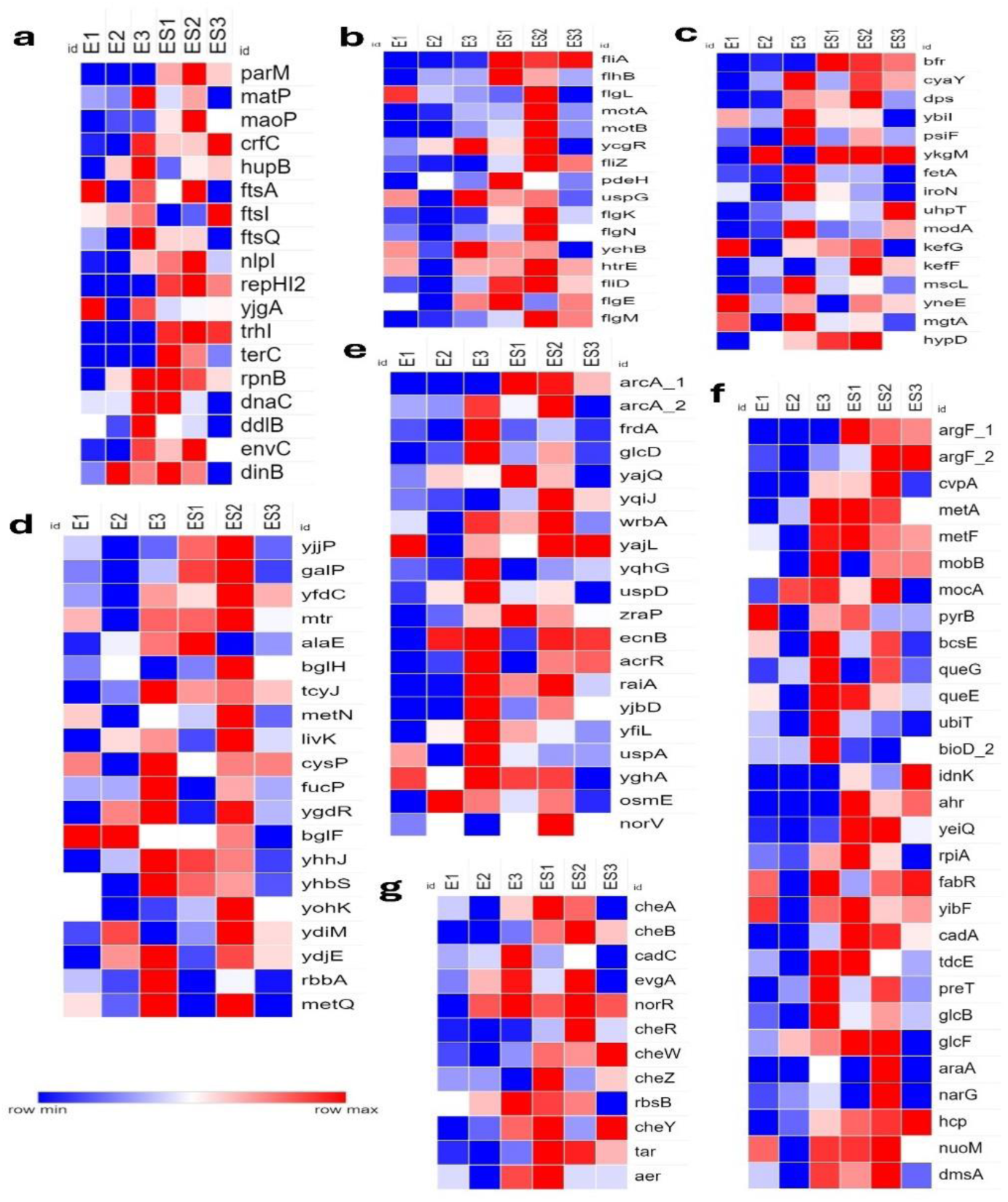
Heat maps of transcriptional responses of *E. coli* 47-1826 in co-culture with *S.* Heidelberg based on functional analysis. (**a**) cell proliferation, (**b**) adherence and bacterial motility, (**c**) ions homeostasis and regulation, (**d**) Outer membrane proteins and transmembrane transporters, (**e**) stress response regulation, (**f**) cellular metabolism, (**g**) signal transduction and chemotaxis. The transformed Fragments/Kilobase of Transcript/Million (FPKM) numbers (normalized read counts) are illustrated in color. Blue indicates low expression, whereas red indicates high expression.

### 2.4. Enriched biological pathways of *S.* Heidelberg 18-9079 in co-culture with *E. coli* 47-1826

Gene enrichment analysis categorized the SDEGs according to their respective biological pathways and metabolic functions to highlight the molecular and cellular processes involved in the interactions between *S*. Heidelberg 18-9079 and *E. coli* 47-1826 when they were co-cultured. Briefly, in *S*. Heidelberg 18-9079, a significant downregulation was observed for the processes involved in cell division septum assembly formation, glycogen biosynthesis, metal ion transport, putrescine catabolism, response to reactive oxygen species and hydrogen peroxide, ATP-independent protein folding, and amino acid catabolism. In contrast, biological processes involved in the metabolism of ribose phosphate, ribonucleotides, nucleoside phosphate, nucleotides, pyrimidine ribonucleotides, nucleoside-monophosphate, nucleobase-containing small molecules, and biosynthesis of cellular nitrogen compounds, nucleotides, and ribonucleotides remained upregulated in *S*. Heidelberg 18-9079 in the presence of *E. coli* 47-1826 (Figure 6).

**Figure 6:**
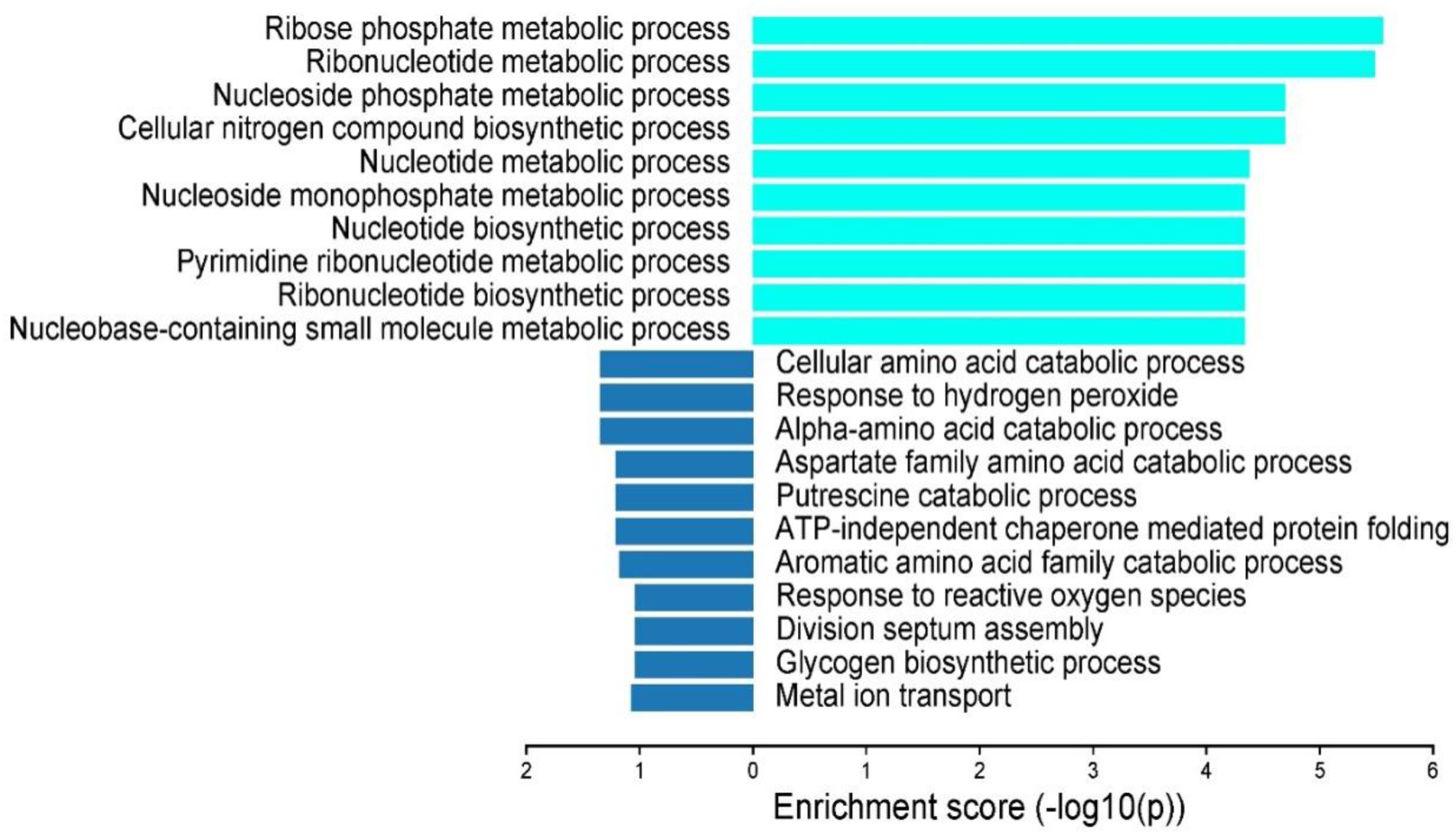
Enriched biological pathways of SDEGs in *S.* Heidelberg 18-9079 in the presence of commensal *E. coli* 47-1826. X-axis shows the enrichment score (-log 10 p-value).

### 2.5. Enriched biological pathways of *E. coli* 47-1826 in co-culture with *S*. Heidelberg 18-9079

In *E. coli* 47-1826, gene enrichment analysis of SDEGs revealed a significant upregulation of the molecular and cellular processes especially involved in the arginine metabolism, arginine catabolism to glutamate and succinate, maturation of protein by iron-sulfur cluster transfer, fermentation, catabolism of glutamate and tryptophan, elevation of intracellular pH, response to L-cysteine, and protein folding. Whereas, biological processes involved in chemotaxis, flagellum-dependent cell motility, aerotaxis, methionine biosynthesis and metabolism, positive regulation of ion transmembrane transporter activity, and thermotaxis were downregulated in *E. coli* 47-1826 in the presence of *S*. Heidelberg 18-9079 (Figure 7).

**Figure 7:**
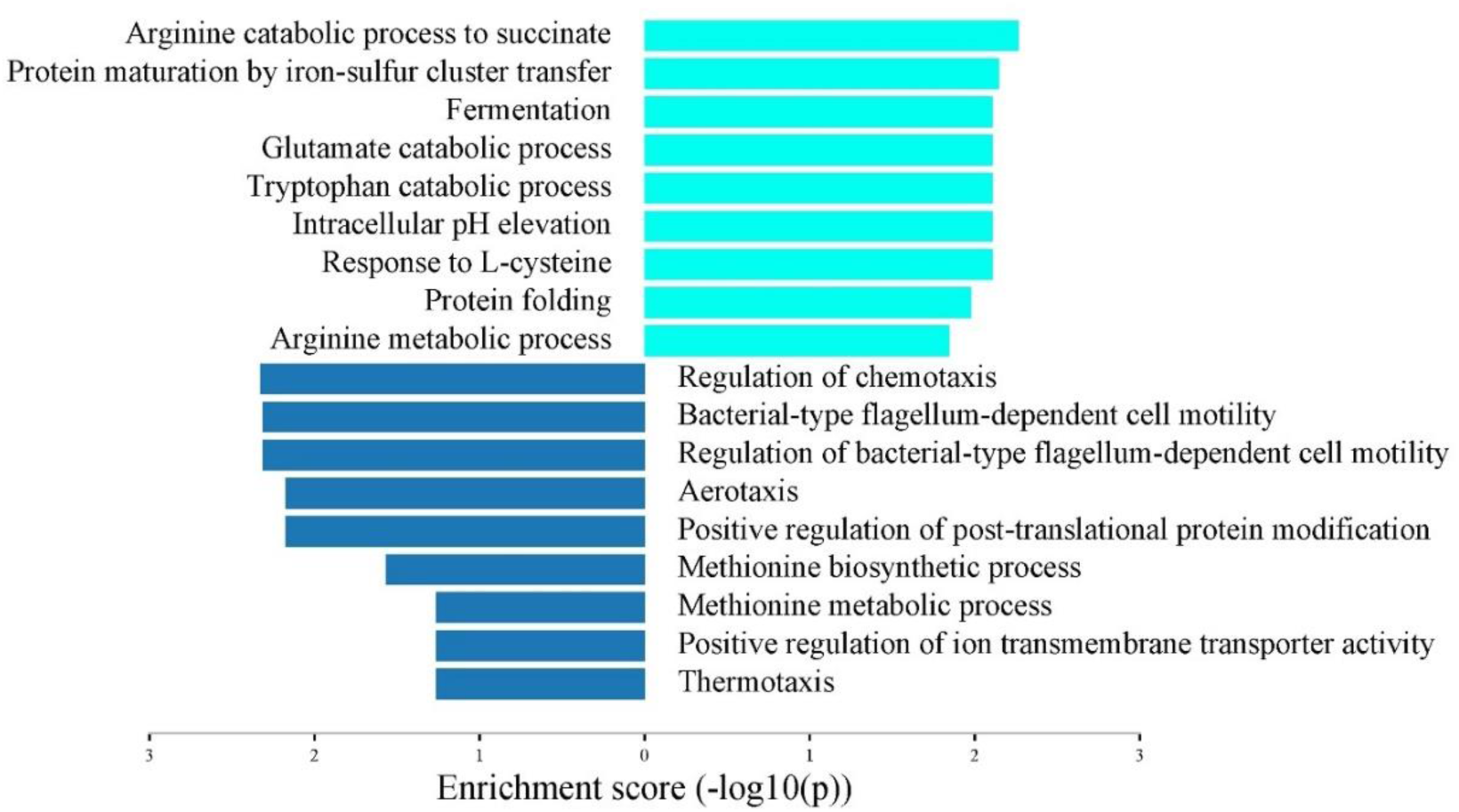
Enriched biological pathways of SDEGs in commensal *E. coli* 47-1826 in the presence of *S.* Heidelberg 18-9079. X-axis shows the enrichment score (-log 10 p-value).

### 2.6. Bacterial growth patterns

When an equal cell number (2.66×10^7^ CFU/mL) of each *S*. Heidelberg 18-9079 and *E. coli* 47-1826 were co-cultured for 24 h, the viable colony count of *S*. Heidelberg 18-9079 (1.63 ± 0.06 log_10_ CFU/mL) in the co-culture was significantly less than the *Salmonella* grown alone (2.2 0 ± 0.07 log_10_ CFU/mL) (p= 0.017). Similarly, the colony count of *S*. Heidelberg 18-9079 (1.63 ± 0.06 log_10_ CFU/mL) in the co-culture was significantly less than the commensal *E. coli* 47-1826 (2.08 ± 0.10 log_10_ CFU/mL) (p = 0.003). However, the difference observed between the colony counts of commensal *E. coli* 47-1826 (2.27 ± 0.03 log_10_ CFU/mL) grown alone and in co-culture (2.08 ± 0.10 log_10_ CFU/mL) (p =0.1013) was statistically insignificant (Figure 1).

### 2.7. Functional annotation of significantly differentially expressed genes of *S.* Heidelberg 18-9079 when co-cultured with commensal *E. coli* 47-1826

#### 2.7.1. Cell proliferation

The results demonstrated significantly low expression of *S*. Heidelberg genes involved in cell proliferation in the presence of commensal *E. coli,* which aligned with the reduced *Salmonella* colony counts (CFU/ml) recorded during the experiment (Figure 1). Specifically, twenty genes, including five genes associated with cell wall organization, cell shape regulation, and cell wall biogenesis (*rlpA, yeah, mtgA, idtA, sylB),* four genes involved in RNA/DNA endonuclease activity (*smrB, rbn, ghoS,* and *gpM-2*), two genes each for cell septum assembly formation during bacterial cell division (z*apA* and *zapB)*, cell division and modulation (*cedA, nlpD),* DNA repair (*recN, yhbO),* and one gene each for protein modification (*sixA)*, small ribosomal biogenesis (*rimP)*, protein folding (*cbpA*), RNA helicase activity (*dbpA*), and cell envelope modulation (*cpxP*) were downregulated in *S.* Heidelberg. Contrary to this observation, two genes associated with the type II toxin-antitoxin system (*ccdA* and *ccdB)* in *S.* Heidelberg were upregulated (Figure 4a).

#### 2.7.2. Pathogenicity and virulence factors

Twelve genes involved in the pathogenicity and virulence of *Salmonella* were downregulated in *S*. Heidelberg. Particularly, the genes involved in the regulation of type III secretion system effectors (TTSS) (*sopE, sopF*), superoxide dismutase (*sodC-2, sodA*), and secretion system apparatus proteins (*ssaD*) were downregulated. Additionally, genes involved in the biosynthesis of polyamines, such as the putrescine precursor/arginine decarboxylase (*speA*), ornithine decarboxylase (*speF*), putrescine transporter (*potF*), and putrescine catabolism enzymes (*patD* and *ygjG*), were suppressed. Furthermore, there was notable downregulation of genes related to flagellar protein biosynthesis (*flhE*) and those involved in the flagellar biosynthesis/type III secretory pathway (*orgA*) (Figure 4b).

#### 2.7.3. Biofilm formation

This study exhibited the expression of genes vital for biofilm formation and regulation in *Salmonella*. A significant downregulation was observed for four genes (*bsmA, bssR, adrA,* and *bapA)* associated with biofilm formation, as well as for two genes (*mlrA* and *mlrB*) that play a role in *csgD* transcription, which is the master regulator of biofilm formation and cellulose biosynthesis. Conversely, an upregulation was observed for *bhsA*, involved in reduced biofilm formation (Figure 4c).

#### 2.7.4. Antimicrobial resistance and multidrug efflux

In this study, several genes in *S.* Heidelberg implicated in AMR and multidrug efflux were downregulated significantly in the presence of commensal *E. coli*. These included the genes that encode an MDR transporter (*mdtM)*, multiple AMR protein (*marB*), macrolide transporter subunit and macrolide ABC transporter ATP-binding proteins/permeases (*macA, macB*), multidrug efflux and toxic compound extrusion (MATE) transporter activation protein (*mdtK*), multidrug efflux pump activation proteins (*mdtK, acrB, mdtG*), and a protein involved in the regulation of the proton-dependent efflux pump (*mdtM*) (Figure 4d). Furthermore, the ESC gene *ctmX1* was also downregulated in *S*. Heidelberg when it was co-cultured with commensal avian *E. coli*.

#### 2.7.5. Ion homeostasis

Fifteen genes associated with metal ion acquisition, utilization, transportation, and storage were downregulated in *S*. Heidelberg. Specifically, five genes involved in iron transport and uptake (*sitA, sitB, sitC, sitD* and *foxR*), three in zinc-ion transmembrane transport (*znuB, znuC* and *zitB*), one each in the iron mobilization/bacterioferritin-associated ferredoxin (*bfd*), zinc-ion binding (*ZinT*), copper homeostasis (*cutC*), cation transportation (*chaB*), and magnesium-ion regulation (*yrbL*) expressed significant suppression in *S.* Heidelberg in addition to the genes involved in divalent metal acquisition and transport (*pspA*) and monoatomic ion channel activity (*ybiO*) (Figure 4e).

#### 2.7.6. Cell metabolism

Several genes involved in critical cellular metabolic processes of *S.* Heidelberg also demonstrated a significant downregulation. These included the genes involved in the catabolism and transport of ethanolamine (*eutA, eutB, eutH,* and *eutJ*), protein biosynthesis and catabolism (*rmf, yidZ, deoC, iscR, caiF, slyA, csiE, mcbR*, *dsbI,* and *htpx*), carbohydrate biosynthesis and gluconeogenesis (*galU, ubiC, sdaB,* and *glgX-2*), amino acid metabolism (*glnQ*, *glnA, hutU, hutH,* and *cdsH),* lipid, lipopolysaccharide, and fatty acid metabolism (*yegS, fadR, fadJ, fadI, fadE, cfa,* and *wzzB,*), trehalose biosynthesis (*otsA, otsB, treY,* and *treZ*) glutathione degradation (*ggt*), and glycan biosynthesis (*glgB*) (Figure 4f).

#### 2.7.7. Outer membrane proteins and transmembrane transporters

The expression of genes associated with outer membrane proteins (OMPs) and transmembrane transport of xenobiotic (*punC*), metalloprotein (*yjdN*), nucleosides (*nupC, impX*), proteins and amino acids (*exbB, exbD*, *glnQ*), sugars (*galP, glpT*), formate (*focA*), lipoprotein localization to the outer membrane (*lolA*), coenzyme transport and metabolism (*yieE*), and outer membrane protein (*ompX*) remained notably downregulated. We also observed a considerably low expression of *smvA*, a gene involved in the transmembrane transportation of acriflavine and other quaternary ammonium compounds (Figure 4g).

#### 2.7.8. Stress response regulation

The expression of genes responsible for stress response mechanisms in *S.* Heidelberg showed significant downregulation. These downregulated genes are specifically involved in the universal stress response (*uspA, uspC,* and *uspE*), SOS response (*yebG, sulA*), glutathione activity (*grxB, yfcG*), and peroxide damage (*yhcN-B*), response to toxic substances (*ecnB*), cellular responses to DNA protection/repair (*ogT*), aldehydes dehydrogenation to mitigate the oxidative/electrophilic stresses (*aldB*), and oxidative stress/phosphorus starvation stress (*psiF*) (Figure 4h).

#### 2.7.9. Signal transduction and chemotaxis

Several genes involved in signaling and chemotaxis in *S.* Heidelberg, particularly those associated with phosphorelay signal transduction and predominantly responding to low levels of external divalent cations zinc and magnesium (*phoP*), phosphorelay sensory kinase/sensory histidine kinase (*narX*), and phosphate regulator induced by phosphate starvation *(phoH),* periplasmic sensors in multi-component regulatory systems/response regulators in the Tor respiratory system (*torT*), and chemotaxis (*mglB*) were suppressed. In contrast, *mgrB*, a negative regulator of the bacterial phosphorelay signal transduction system, was upregulated (Figure 4i).

### 2.8. Functional annotation of significantly differentially expressed genes of commensal *E. coli* 47-1826 when co-cultured with *S.* Heidelberg 18-9079

#### 2.8.1. Cell proliferation

Our results showed a significantly high expression of *E. coli* genes involved in cell proliferation in the presence of *Salmonella,* consistent with the observed increase in *E. coli* colony counts during the experiment (Figure 1). These 18 upregulated *E. coli* genes included five genes associated with chromosome segregation, organization and condensation (*parM*, *matP*, *moaP*, *crfC*, *hupB*), three genes involved in the barrier septum assembly formation (*ftsA, ftsI, ftsQ*), three genes for cell division (*nlpl*, *repHI2, yjgA*), and one gene each involved in 3-5 prime DNA helicase activity and unwinding of duplex DNA (*trhL*), protection from abnormal sticking together or degradation of chromosomes (*terC*), double-stranded DNA endonuclease activity and DNA recombination (*rpnB*), elongation of DNA-strand during DNA replication (*dnaC*), cell shape regulation (*ddIB*), cell separation after cytokinesis (*envC*), and DNA-dependent DNA replication (*dinB*) (Figure 5a).

#### 2.8.2. Adherence and bacterial motility

The results indicated an upregulation of 16 genes critical for the adherence and bacterial motility in *E. coli* in the co-culture. These included six genes responsible for bacterial-type flagellum-dependent cell motility (*fliA*, *flhB, flgL, motA, motB, ycgR*), three genes for regulation of bacterial-type flagellum-dependent cell motility (*fliZ, pdeH, uspG*), two genes each for the formation of bacterial-type flagellum assembly (*flgK, flgN*), and pilus assembly formation and cell adhesion (*yehB, htrE*). One gene each for bacterial-type flagellum-dependent cell motility and cell adhesion (*fliD*), bacterial-type flagellum-dependent swarming motility (*flgE*), and bacterial-type flagellum organization (*flgM*) was also upregulated (Figure 5b).

#### 2.8.3. Ion homeostasis

In this study, 17 genes associated with metal ions acquisition, storage, and homeostasis were upregulated in commensal *E. coli* grown together with S. Heidelberg. Particularly, genes responsible for intracellular sequestration of iron-ions (*bfr*), iron-sulfur cluster assembly formation (*cyaY*), cellular iron-ions homeostasis (*dps*), iron-ions binding (*hypD*), zinc-ion binding (*ybiI*) cellular response to phosphate starvation (*psiF*), and cellular response to zinc-ion starvation (*ykgM*) were upregulated. Genes associated with transmembrane movement and transportation of iron (*fetA*), siderophores (*iroN*), phosphate (*uhpT*), molybdate-ion (*modA*), potassium (*kefG, kefF*), monoatomic-ions (*mscL*), chloride (*yneE*), and magnesium (*mgtA*) were also significantly upregulated in the commensal *E. coli* in co-culture with *S*. Heidelberg (Figure 5c).

#### 2.8.4. Signal transduction and chemotaxis

Several genes involved in signaling and chemotaxis in commensal *E. coli*, especially five genes associated with bacterial phosphorelay signal transduction (*cheA, cheB, cadC, evgA, norR,*), four genes with chemotaxis and regulation of chemotaxis (*cheR, cheW, cheZ*, *rbsB*), and one gene each for signal-complex assembly formation (*tar*), thermotaxis (*cheY*) and positive aerotaxis (*aer*) were significantly upregulated (Figure 5g).

#### 2.8.5. Stress response regulation

Twenty genes responsible for stress response mechanisms in commensal *E. coli* were upregulated. Importantly, genes involved in aerobic respiration control (*arcA-1* and *arcA-2*), cellular response to DNA damage stimulus (*frdA, glcD, yajQ*, *yqiJ*), oxidative stress (*wrbA*, *yajL*, *yqhG*), superoxides (*uspD*), cellular response to cell-envelope stress (*zraP*), toxic substances (*ecnB*), and xenobiotic stimulus (*AcrR*). Furthermore, genes associated with bacterial dormancy process in unfavorable conditions (*raiA*), general and universal stress responses (*yjbD, yfiL*, *uspA*), osmotic stress (*yghA*, *osmE*), and response to nitric oxide (*norV*) were also notably upregulated in commensal *E. coli* in the presence of *S*. Heidelberg (Figure 5e).

#### 2.8.6. Outer membrane proteins and transmembrane transporters

The expression of genes associated with OMPs and vital for transmembrane transport of succinate (*yjjP*), galactose (*galP*), formate (*yfdC*), amino acids (*mtr, alaE*), polysaccharide (*bglH*), cysteine (*tcyJ*), D-methionine (*metN*, *metQ*), leucine (*livK*), thiosulfate (*cysP*), fucose (*fucP*), histidne (*ygdR*), and trehalose (*bglF*) were upregulated in *E. coli*. Furthermore, genes involved in ABC-type transportation activity (*rbbA*, *yhhJ*, *yhbS*) and 3-hydroxypropanoate export (*yohK*) were also significantly upregulated. We also noted the upregulation of genes (*ydiM, ydjE*) associated with the major facilitator superfamily (MFS) type transmembrane transport activity (required for cellular Fe and Cu homeostasis for the biogenesis of different heme-Cu oxygen reductases,) in the commensal *E. coli* (Figure 5d).

#### 2.8.7. Cell metabolism

Several genes involved in critical cellular metabolic processes were significantly upregulated in *E. coli*. These genes included those associated with the biosynthesis of arginine (*argF-1* and *argF-2*), toxin production (*cvpA*), methionine synthesis (*metA* and *metF*), Mo-molybdopterin cofactor biosynthetic process (*mobB* and *mocA*), pyrimidine metabolism (*pyrB*), cellulose production (*bcsE*), queuosine synthesis (*queG* and *queE*), ubiquinone production (*ubiT*), and biotin synthesis (*bioD-2*). Similarly, genes involved in the metabolism of D-gluconate (*idnK*, *ahr*), mannitol (*yeiQ*), ribose (*rpiA*), fatty acids (*fabR*), and glutathione (*yibF*) were also upregulated. Furthermore, several other genes responsible for catabolism, particular those related to lysine (*cadA*), L-threonine to propionate (*tdcE*), thyamine (*preT*), glyoxylate (*glcB* and *glcF*), and arabinose (*araA*), were similarly upregulated. Notably, we observed a significant upregulation of genes involved in nitrate assimilation (*narG*), cellular oxidant detoxification (*hcp*), as well as aerobic (*nuoM*) and anaerobic respiration (*dmsA*) in commensal *E. coli* co-cultured with *S*. Heidelberg (Figure 5f).

### 2.9. Validation of gene expression data

Real-time quantitative PCR (RT-qPCR) was employed to validate the reliability and accuracy of RNA sequencing data by quantifying relative expression patterns of six selected genes in *S*. Heidelberg 18-9079 and two genes in *E. coli* 47-1826 under the same co-culture conditions used in the transcriptome analysis experiment. Consistant with the RNAseq analysis, RT-PCR demonstrated a log_2_ fold-change (FC) expression of *sitA* (−0.9108), *punC* (−0.2713), *bsmA* (−0.8899), *mlrA* (−32.09), *sopE* (−0.117), and *bhsA* (2.1006) in *S*. Heidelberg 18-9079, and the log_2_ FC expression of *repHI2* (14.206) and *arcA* (3.499) in *E. coli* 47-1826, thereby validating our transcriptomic profiling data.

## 3. Discussion

Our study focused on determining the differential gene expression in antibiotic-resistant *S.* Heidelberg 18-9079 in the presence of commensal *E. coli* 47-1826 to delineate the effect of commensal *E. coli* on fitness, virulence, and AMR in antibiotic-resistant nontyphoidal *Salmonella* with a long-term goal of developing pre-harvest intervention strategies to reduce *Salmonella* colonization of poultry thereby reducing foodborne salmonellosis in humans.

The results of gene enrichment analysis indicate the downregulation of genes involved in the formation of cell-division septum assembly in *S*. Heidelberg in the presence of commensal *E. coli.* Cell septum is a critical component for proper cell division and bacterial propagation in hosts (Willis & Huang, 2017). In this context, the downregulation of genes (*zapA* and *zapB*) in *S*. Heidelberg indicates a loose integration of cell septum assembly. This leads to malfunctions or a complete absence of the z-ring protein (FtsZ), resulting in either a complete halt in cell division or disruptions in the regulatory processes that initiate and complete bacterial cell division (Ebersbach et al., 2008; Fernández-Fernández et al., 2021; Misra et al., 2018). Furthermore, the overexpression of genes (*ccdA*, *ccdB*) involved in post-segregation killing of bacterial cells may be associated with impaired cell division in *Salmonella* due to low expression of *zapA* and *ZapB* genes. The expression of these genes (*zapA*, *ZapB* and *ccdA*, *ccdB*) observed in this study could be critical for inducing cell death and reduced growth pattern of *Salmonella* in co-culture (Fraikin et al., 2020; Tripathi et al., 2012, Fernández-Fernández et al., 2021). Additionally, reduced expression of cell division activator (*cedA*) and cell damage protection protein (*nlpD*) may further suppress cell division and growth of *S.* Heidelberg in response to stress elicited by commensal *E. coli* (Burin et al., 2014; Finn et al., 2013). Therefore, the disruption of tightly and precisely controlled interplay of coordinated complex cellular processes by commensal *E. coli* involved in *S.* Heidelberg multiplication and growth may lead to reduced multiplication of *Salmonella* in the chicken gastrointestinal tract which is essential for its colonization and persistence in chickens, as well as for its transmission to new hosts (Nair et al., 2021; Willis & Huang, 2017).

This study provided valuable insights into polyamine biosynthesis and putrescine catabolism, the pathways crucial for *Salmonella* virulence (Jelsbak et al., 2012; Nair et al., 2023; Schroll et al., 2014). Suppression of putrescine catabolism results in intracellular putrescine accumulation, critically inhibiting the synthesis of spermidine and spermine, which in turn may compromise the virulence, motility, cell surface adhesion, protein synthesis, and viability of *Salmonella* (Espinel et al., 2016; Guerra et al., 2020; Nair et al. 2024, 2023). In this study, the expression of genes involved in putrescine biosynthesis from arginine (*speA)* or ornithine (*speF*) remained downregulated, while the *speE* involved in putrescine catabolism into spermidine also downregulated (Jelsbak et al., 2012; Schroll et al., 2014). The reduced expression of the compensatory putrescine transporter (*PotF*) of *S.* Heidelberg may further diminish its virulent potential (Turnbough & Switzer, 2008). Polyamine depletion affects the expression of genes associated with virulence, pathogenicity, and stress responses in *Salmonella.* Therefore, the downregulation of polyamine biosynthesis observed in this study could be critical to reduce the virulence and pathogenicity of *S*. Heidelberg by suppressing the type III secretion system (*sopE* and *sopF*) and the effector protein (*ssaD*) in *S*. Heidelberg (Guerra et al., 2020). Therefore, the loss of polyamine biosynthesis and transport ability can substantially limit the pathogenicity of *Salmonella* by lowering the expression of virulence genes located on pathogenicity islands (Guerra et al., 2020; Nair et al., 2023). The lower expression of flagellar biosynthesis (*fhlE*) might diminish the motility of *S.* Heidelberg, which is essential for colonization and tissue invasion, thereby compromising *Salmonella* survival and virulence. Reduced *orgA* expression might further reduce *S*. Heidelberg mobility and invasiveness (Lee et al., 2015; Nair et al., 2023; Russell et al., 2004). Furthermore, gene *fadR* involved in the biosynthesis of unsaturated fatty acids is vital to maintain a well-balanced ratio of unsaturated and saturated fatty acids for sustained biophysical properties of cell membrane phospholipids essential for bacterial growth and survival, probably through biofilm formation. Therefore, a significantly reduced expression of *fadR* gene might further compromise the growth and survival of *Salmonella* (Herman et al., 2022).

Likewise, enrichment analysis also provided considerable insight into metal ion acquisition, transportation, and utilization essential for the survival, pathogenicity, growth, and persistence of *Salmonella* inside the host (Battistoni et al., 2017; Huang et al., 2018; Mey et al., 2021; O’connor et al., 2006; Samanovic et al., 2012). It has previously been reported that interactions with commensal *E. coli* suppress the expression of genes involved in metal ion homeostasis in *Salmonella* (Deriu et al., 2013; Nairz et al., 2018). In this context, the downregulation of the inner membrane siderophore-independent system ABC-type transporter genes (*sitA*, *sitB, sitC*, and *sitD)* responsible for iron acquisition and transportation suggests reduced iron homeostasis in *Salmonella,* which is crucial for its virulence and persistence (Deriu et al., 2013; Mey et al., 2021). Furthermore, suppression of the transcriptional regulator of the xenosiderophore ferrioxamine transporter (*foxA*) in conjunction with lower expression of xenosiderophore-mediated iron uptake (*foxR)* might further diminish the iron uptake by *Salmonella* (Saldaña-Ahuactzi & Knodler, 2022). Similarly, downregulation of *znuB*, *znuC*, *zniT,* and *zitB* would compromise the ability of *Salmonella* to maintain free intracellular zinc levels required for normal cellular homeostasis. The lack of expression of *ZnuB*, *ZnuC,* and *ZnuA* can also noticeably reduce the ability of *Salmonella* to adapt to its various environments and pathogenicity (Battistoni et al., 2017; Huang et al., 2018). Moreover, suppression of genes responsible for copper and magnesium ion homeostasis may render a variety of enzymes critical for normal cellular function, cell growth, differentiation, and survival of *Salmonella* inactive (O’connor et al., 2006; Samanovic et al., 2012).

This study also sheds light on the expression of genes essential for response regulation against reactive oxygen species (ROS). Lethal stress leads to intracellular accumulation of ROS and induces oxidative stress, damaging essential macromolecules. However, bacteria employ numerous defensive proteins to detoxify ROS and counteract the damage incurred by ROS (Rhen, 2019; Zhao & Drlica, 2014). Therefore, the observed suppression of genes encoding superoxide dismutase (*sodC-2*) might subject *S*. Heidelberg to oxidative stress and impaired colonization (Ammendola et al., 2005; Pacello et al., 2008). In conjunction with this, suppression of another superoxide dismutase protein encoded by *sodA* may further increase the vulnerability of *S*. Heidelberg to ROS, leading to low growth and reduced survivability (Wang et al., 2018). Downregulation of another protein OmpX, critical in responding to ROS stress, may additionally enhance the sensitivity of *S*. Heidelberg to hydrogen peroxide stress in the presence of commensal *E. coli* (Briones et al., 2022). Similarly, a notable reduction in the expression of universal stress proteins (*UspA, UspC,* and *UspE*) may further reduce the potential of *Salmonella* to withstand and survive under stressful conditions (Liu et al., 2007; O’connor et al., 2006). The ability of commensal *E. coli* to compete for luminal oxygen in the gastrointestinal tract also plays a role in reducing intestinal colonization of *Salmonella* (Litvak et al., 2019).

*ArcA*, sometimes touted as a global regulator of gene expression, is an integral part of the two-component system ArcAB (anoxic redox control or aerobic respiration control) in *E. coli* (Brown et al., 2022). Under anaerobic conditions, *ArcA* acts as a global response regulator controlling many operons and regulating diverse metabolic pathways either directly or indirectly Nochino et al., 2020). Therefore, *ArcA* plays a critical role in the survival and growth of *E. coli* in stressful and resource-competitive environments by regulating energy production via metabolic modifications, effective stress response, redox balance, and the acquisition and utilization of several key elements such as nitrogen, amino acids, and iron (Brown et al., 2022). In the current study, alteration in the transcriptomic profile of *S*. Heidelberg in co-culture may be linked to the observed upregulation of *arcA* in commensal *E. coli*. Under anaerobic conditions, *arcA* plays a critical role in balancing catabolic efficiency (energy production) and anabolic processes (biomass growth) through global regulation of metabolism. Our findings show a significantly higher growth of commensal *E. coli* 47-1826 in the co-culture along with a significant downregulation of genes critical for growth, persistence, and antimicrobial resistance dissemination observed in *S*. Heidelberg 18-9079. This downregulation could be associated with the elevated expression of the *arcA* in the commensal *E. coli*, which might allow *E. coli* to competitively exclude *S*. Heidelberg from the chicken intestines (Loui et al., 2022). The expression of *arcA* in conjunction with other stress stimulus leads to the metabolic reprogramming in *E. coli*, potentially enhancing the survival and growth of commensal *E. coli* under competitive conditions (Brown et al., 2022, 2023). However, the connotation that the higher expression of *arcA* in commensal *E. coli* 47-1826 downregulated the expression of genes critical for colonization, pathogenicity, and survival in *S*. Heidelberg is speculative at this point, and further research is needed to substantiate such an association. Overall, the exact mechanism behind the transcriptomic shift occurred in *S*. Heidelberg 18-9079 in the presence of a commensal *E. coli* 47-1826 strain remains unclear.

Commensal *E. coli* is one of the earliest colonizers of poultry intestines and predominates the gut microbiota population at a very young age (Musa et al., 2021). Sufficient intestinal colonization is essential for the effectiveness of probiotic bacteria. From this perspective, intestinal commensal *E. coli* has a competitive edge over other bacteria. The ability to naturally colonize chicken intestines exempts commensal *E. coli* 47-1826 from the multiple stresses encountered by other bacteria while passing through different intestinal segments to attain sufficient colonization in the chicken gut (Nair et al., 2021; Patterson & Burkholder, 2003). In addition, a significant downregulation of genes critical for metabolic and cellular pathways conferring fitness, virulence, and AMR dissemination potential in *S*. Heidelberg provides evidence that commensal *E. coli* 47-1826 as probiotic or its products as prebiotics can potentially be used to mitigate drug-resistant *S*. Heidelberg and other nontyphoidal *Salmonella* colonization and fecal shedding at the preharvest level in the poultry production system (Deriu et al., 2013; Sun et al., 2022; Wu et al., 2023).

This study offers valuable insights into the potential role of commensal *E. coli* in controlling nontyphoidal *Salmonella* in poultry, as well as other animal species, at the farm level. However, there are a few limitations of the study to consider. First, this study evaluated only one strain of poultry commensal *E. coli* and one strain of antibiotic-resistant *S.* Heidelber*g*; thus, the transcriptomic changes may not be valid for other strains of commensal *E. coli* and *Salmonella*. Second, since this study was an in vitro experiment, and therefore, the results need to be confirmed through in-vivo experiments or field trials. Despite these limitations, the results of this study shed light on the behavior of *S.* Heidelberg in the presence of commensal *E. coli* 47-1826 as revealed through differential transcriptomics profiling. This study also provides new insights into the potential ability of commensal *E. coli* 47-1826 to prevent colonization of nontyphoidal *Salmonella* in poultry and offers the framework to explore the mechanisms by which commensal *E. coli* downregulates the expression of genes critical for fitness, virulence, and AMR potential in *S.* Heidelberg. Finally, this study highlights the promise of using species-specific commensal strains of *E. coli* to avert the colonization potential of *Salmonella* in poultry, and possibly in other animal species and humans.

## Conclusion

Our research demonstrated that the commensal *E. coli* strain 47-1826 suppresses the growth of the ESC-resistant *S.* Heidelberg strain 18-9079. Additionally, it effectively downregulates the expression of key genes responsible for critical metabolic pathways and cellular processes that drive the propagation, survival, pathogenicity and virulence, and AMR of *S*. Heidelberg. Specifically, the expression of genes associated with cell proliferation, pathogenicity and virulence, biofilm formation, AMR and drug efflux, ion hemostasis and regulation, signal transduction and chemotaxis, stress response management, outer membrane proteins and transmembrane transportation, and cellular metabolism was significantly downregulated. Whereas, in the commensal *E. coli* strain 47-1826 the genes involved in the cell proliferation, bacterial adherence and motility, ion homeostasis, signal transduction and chemotaxis, stress response to xenobiotics, outer membrane proteins and transmembrane transportation, and vital cellular metabolism were significantly upregulated. Notably, the upregulation of *arcA* might have enabled the commensal *E. coli* 47-1826 strain to effectively compete, survive and grow in the co-culture with *S.* Heidelberg 18-9079. Given these compelling findings, we conclude that commensal *E. coli* significantly downregulated critical cellular pathways and functions that contribute to the fitness, virulence, and AMR to *S.* Heidelberg 18-9079, which play a crucial role in the colonization of *S*. Heidelberg in the chicken gut. Therefore, commensal *E. coli* 47-1826 or its products could be used as an alternative to antibiotics for controlling the colonization of antibiotic-resistant *S.* Heidelberg in poultry.

## 4. Material and Methods

### 4.1. Bacterial strains

*S*. Heidelberg 18-9079 strain isolated from the liver of a turkey, which is resistant to ESC and harboring *bla*_CTX-M1_ (Denagamage et al., 2019), and the commensal strain of *E. coli* 47-1826 isolated from the intestines of a healthy broiler chicken were used in co-culture studies.

### 4.2. Bacterial culture

*S*. Heidelberg 18-9079 and *E. coli* 47-1826 were co-cultured in Luria Bertani (LB) broth (BD, Difco, Franklin Lakes, NJ, USA) at an equal concentration (2.66 × 10^7^ CFU/mL, OD_600_ = 0.01) (Spectrophotometer, Thermo Fischer Scientific, USA) and incubated for 24 h at 37°C under anaerobic conditions, created with AnaeroPack-Anaero (Thermo Scientific, Swedesboro, NJ), with shaking. The *S.* Heidelberg 18-9079 and *E. coli* 47-1826 were also cultured separately under similar concentrations to serve as controls. For each bacterial culture, three biological replicates were processed for RNA extraction.

### 4.3. RNA extraction and rRNA depletion

Total RNA (RNA integrity number, RIN ≥ 8.5) was extracted using the RiboPure RNA Purification Kit (Invitrogen, Waltham, MA, USA) and further processed using a MICROBExpress Bacterial mRNA Enrichment Kit (Invitrogen, Waltham, MA, USA) to enrich for mRNA by depleting ribosomal RNA (rRNA) according to manufacturer’s instructions. The quality and quantity of purified mRNA were determined using a NanoDrop™ 2000/2000c spectrophotometer (Thermo Fisher Scientific, Wilmington, DE, USA). The quality of total RNA and mRNA were further assessed by 1% agarose and 1% denaturing formaldehyde gel electrophoresis, respectively.

### 4.4. Library construction and RNA sequencing

Libraries were prepared using the TruSeq^®^ Stranded mRNA Library Kit (Illumina Inc., San Diego, CA, USA). The enriched mRNA was fragmented and primed to synthesize first- and second-strand cDNA. The double-stranded cDNA fragments were adenylated at the 3’ end, ligated to multiple index adapters (TruSeq RNA Combinatorial Dual Indexes; Illumina Inc., San Diego, CA, USA), and enriched to amplify the amount of cDNA in the library. Finally, cDNA libraries were multiplexed and clustered in one lane of a flow cell for sequencing on a MiSeq® System (Illumina, San Diego, CA, U.S.A.) with 2 × 300 bp paired-end (PE) reads and 100 M reads coverage. *S.* Heidelberg 18-9079 and commensal *E. coli* 47-1826 grown separately at 37°C for 24 h in LB broth were used as controls to determine differential gene expressions. All sequences generated were deposited in the NCBI database under accession GSE 276976.

### 4.5. Differential gene expression analysis

Quality assessment, base trimming, normalization, read mapping, and differential gene expression analysis were conducted through the CLC Genomics Workbench version 23.0.4 (https://digitalinsights.qiagen.com) (QIAGEN Inc., Redwood City, CA, USA). The raw read sequences were imported into the CLC Genomics Workbench and aligned with *S.* Heidelberg 18-9079 chromosome (NCBI Refseq assembly: GCF_016452005.1) and its ESC-resistance encoding plasmid (NCBI Refseq assembly: NZ _MW 349106.1), and *E. coli* APEC O1 (NCBI Refseq assembly: GCF_000014845.1). Gene expressions were calculated using RPKM (reads/kilobase of exon model/million mapped reads) and applying the equation RPKM = number of gene reads/mapped reads (millions) × gene length (kb). Significant variations in gene expression (upregulation or downregulation) were determined after employing the false discovery rate (FDR) (Pawitan et al., 2005). The TMM normalization (trimmed mean of M values) described for whole transcriptome RNAseq technology was applied to determine differential gene expression (DGE) in the two groups (*S.* Heidelberg 18-9079 + *E. coli* 47-1826 vs *S.* Heidelberg 18-9079, and *S.* Heidelberg 18-9079 + *E. coli* 47-1826 vs *E. coli* 47-1826). The results of samples submitted in triplicate were averaged to determine the fold change (FC) in gene expression. Filtration criteria, such as FDR ≤ 0.05, FC ≥ 2.0, p-value ≤ 0.01, and FDR ≤ 0.05, FC < 1.0, p-value ≤ 0.01 were employed as cutoff values for DGE for upregulation and downregulation, respectively.

### 4.6. Bacterial growth pattern

*S*. Heidelberg 18-9079 and *E. coli* 47-1826 were cultured separately and in co-culture at an equal concentration (OD_600_ = 0.01) in 50 ml tubes containing LB broth and incubated anaerobically at 37°C for 24 h with shaking. Subsequently, 100-µL volumes of ten-fold serial dilutions (10^-6^) of cultures were plated on McConkey agar and incubated anaerobically at 37°C for 24 h. The number of bacteria or colony-forming units per mL (CFU/mL) were counted to determine the bacterial numbers.

### 4.7. Validation of differential gene expressions

Relative gene expression results of *S*. Heidelberg 18-9079 and commensal *E. coli* 47-1826 grown in co-culture were validated by quantitative real-time PCR (qRT-PCR) in comparison to their respective controls. Six selected genes involved in metal-ion transport, iron acquisition and utilization (*sitA*), detoxification of xenobiotics (*punC*), bacterial survival and biofilm formation (*bsmA* and *bhsA*), motility (*mlrA*), and pathogenicity (*sopE*) in *S*. Heidelberg 189079, and two genes involved in replication (*repHI2*) and aerobic respiration control (*arcA*) in *E. coli* 47-1826 were used in qRT-PCR. The primers used are listed in the supplemental file (S1). The 16S rRNA gene served as an endogenous control. Relative gene expression was determined according to the comparative critical threshold (Ct) method using QuantStudio™ Real-time PCR software V 1.5.2 (Applied Biosystems, Carlsbad, CA, USA). Finally, the data were normalized to the endogenous control 16S rRNA to determine the expression of selected genes in co-cultured bacteria compared to single bacterial cultures.

### 4.8. Statistical analysis, software, and data preparation

All quantitative assays were conducted in triplicates in independent experiments. Differential gene expressions were statistically analyzed using the CLC Genomics Workbench version 23.0.4 after applying the General Linear Model with the negative binomial distribution. A p ≤ 0.01 was considered statistically significant. Gene enrichment analysis was conducted in iDEP 2.0 (http://bioinformatics.sdstate.edu/idep/) to determine different biological pathways governed by significantly differentially expressed genes (SDEGs). Gene enrichment analysis plots were created using SR plot (https://www.bioinformatics.com.cn/), whereas the heat maps were generated in the iDEP 2.0 and Morpheus (https://software.broadinstitute.org/morpheus/) by applying Euclidean distance with average linkage clustering for iDEP 2.0. Antibiotic-resistant genes were identified through Comprehensive Antibiotic Resistance Database (https://card.mcmaster.ca/.), while virulence genes were identified using the Virulence Factor of Pathogenic Bacteria database (http://www.mgc.ac.cn/VFs/). The paired t-test was used to analyze the viable bacterial colony counts of *S*. Heidelberg 18-9079 and *E. coli* 47-1826 in co-culture with bacterial colony counts of their respective cultures grown separately. Whereas bacterial colony counts of *S*. Heidelberg 18-9079 and *E. coli* 47-1826 grown in co-cultures were analyzed by unpaired t-test in GraphPad prism (10.2.3). A p ≤ 0.05 was considered significant.

### 4.9. Data availability

Transcriptomic profiles were deposited in Gene Expression Omnibus (GEO) database in the NCBI under accession GSE 276976.

## Supplementary material

Supplementary data are provided in the attached files.

## Ethical statement

This experiment did not require animals; therefore, an ethical statement was not required.

## Acknowledgments

“This research received no specific grant from any funding agency in the public, commercial, or non-profit sectors”. However, the first author (Yasir R. Khan) is a PhD student under the scholarship “US-Pakistan Knowledge Corridor” with partial funding from the College of Veterinary Medicine, University of Florida, United States. We acknowledge Linda Archer for laboratory support during this experiment.

## Conflicts of interest

Authors have no conflicts of interest to declare.

## References

1. Ammendola, S., Ajello, M., Pasquali, P., Kroll, J. S., Langford, P. R., Rotilio, G., Valenti, P., & Battistoni, A. (2005). Differential contribution of sodC1 and sodC2 to intracellular survival and pathogenicity of Salmonella enterica serovar Choleraesuis. Microbes and Infection, 7(4), 698–707. 10.1016/j.micinf.2005.01.005.

2. Battistoni, A., Ammendola, S., Chiancone, E., & Ilari, A. (2017). A novel antimicrobial approach based on the inhibition of zinc uptake in Salmonella enterica. Future Medicinal Chemistry, 9(9), 899–910. 10.4155/fmc-2017-0042.

3. Briones, A. C., Lorca, D., Cofre, A., Cabezas, C. E., Krüger, G. I., Pardo-Esté, C., Baquedano, M. S., Salinas, C. R., Espinoza, M., Castro-Severyn, J., Remonsellez, F., Hidalgo, A. A., Morales, E. H., & Saavedra, C. P. (2022). Genetic regulation of the ompX porin of Salmonella Typhimurium in response to hydrogen peroxide stress. Biological Research, 55(1). 10.1186/s40659-022-00377-3.

4. Brown AN, Anderson MT, Smith SN, Bachman MA, Mobley HLT (2023). Conserved metabolic regulator ArcA responds to oxygen availability, iron limitation, and cell envelope perturbations during bacteremia. mBio14:e01448–23. 10.1128/mbio.01448-23.

5. Brown, A.N., Anderson, M.T., Bachman, M.A. and Mobley, H.L., (2022). The ArcAB two-component system: function in metabolism, redox control, and infection. Microbiology and Molecular Biology Reviews, 86(2), pp.e00110–21.10.1128/mmbr.00110-21

6. Burin, R. C. K., Silva, A., & Nero, L. A. (2014). Influence of lactic acid and acetic acid on Salmonella spp. growth and expression of acid tolerance-related genes. Food Research International, 64, 726–732. 10.1016/j.foodres.2014.08.019.

7. CDC. National Antimicrobial Resistance Monitoring System (NARMS) Now: Human Data. Atlanta, Georgia: U.S. Department of Health and Human Services, CDC. 09/25/2024. www.cdc.gov/narmsnow. Accessed 9/26/2024.

8. Chen, C., Li, J., Zhang, H., Xie, Y., Xiong, L., Liu, H., & Wang, F. (2020). Effects of a probiotic on the growth performance, intestinal flora, and immune function of chicks infected with Salmonella pullorum. Poultry Science, 99(11), 5316–5323. 10.1016/j.psj.2020.07.017.

9. Denagamage, T. N., Wallner-Pendleton, E., Jayarao, B. M., Xiaoli, L., Dudley, E. G., Wolfgang, D., & Kariyawasam, S. (2019). Detection of CTX-M-1 extended-spectrum beta-lactamase among ceftiofur-resistant Salmonella enterica clinical isolates of poultry. Journal of Veterinary Diagnostic Investigation, 31(5), 681–687. 10.1177/1040638719864384.

10. Deriu, E., Liu, J. Z., Pezeshki, M., Edwards, R. A., Ochoa, R. J., Contreras, H., Libby, S. J., Fang, F. C., & Raffatellu, M. (2013). Probiotic bacteria reduce Salmonella typhimurium intestinal colonization by competing for iron. Cell Host and Microbe, 14(1), 26–37. 10.1016/j.chom.2013.06.007.

11. Dewi, G., Nair, D. V. T., Peichel, C., Johnson, T. J., Noll, S., & Kollanoor Johny, A. (2021). Effect of lemongrass essential oil against multidrug-resistant Salmonella Heidelberg and its attachment to chicken skin and meat. Poultry Science, 100(7). 10.1016/j.psj.2021.101116.

12. Ebersbach, G., Galli, E., Møller-Jensen, J., Löwe, J., & Gerdes, K. (2008). Novel coiled-coil cell division factor ZapB stimulates Z ring assembly and cell division. Molecular Microbiology, 68(3), 720–735. 10.1111/j.1365-2958.2008.06190.x.

13. Edison, L. K., Kudva, I. T., & Kariyawasam, S. (2023). Comparative Transcriptome Analysis of Shiga Toxin-Producing Escherichia coli O157: H7 on Bovine Rectoanal Junction Cells and Human Colonic Epithelial Cells during Initial Adherence. Microorganisms, 11(10), 2562. 10.3390/microorganisms11102562

14. Espinel, I. C., Guerra, P. R., & Jelsbak, L. (2016). Multiple roles of putrescine and spermidine in stress resistance and virulence of Salmonella enterica serovar Typhimurium. Microbial Pathogenesis, 95, 117–123. 10.1016/j.micpath.2016.03.008.

15. Fagbamila, I. O., Ramon, E., Lettini, A. A., Muhammad, M., Longo, A., Antonello, K., … & Barco, L. (2023). Assessing the mechanisms of multi-drug resistant non-typhoidal Salmonella (NTS) serovars isolated from layer chicken farms in Nigeria. Plos one, 18(9), e0290754. 10.1371/journal.pone.0290754

16. Fernández-Fernández, R., Hernández, S. B., Puerta-Fernández, E., Sánchez-Romero, M. A., Urdaneta, V., & Casadesús, J. (2021). Evidence for Involvement of the Salmonella enterica Z-Ring Assembly Factors ZapA and ZapB in Resistance to Bile. Frontiers in Microbiology, 12. 10.3389/fmicb.2021.647305.

17. Finn, S., Händler, K., Condell, O., Colgan, A., Cooney, S., McClure, P., Amézquita, A., Hinton, J. C. D., & Fanning, S. (2013). ProP is required for the survival of desiccated Salmonella enterica serovar typhimurium cells on a stainless-steel surface. Applied and Environmental Microbiology, 79(14), 4376–4384. 10.1128/AEM.00515-13.

18. Fraikin, N., Goormaghtigh, F., & Melderen, L. Van. (2020). Type II Toxin-Antitoxin Systems: Evolution and Revolutions. 10.1128/JB.

19. Guerra, P. R., Liu, G., Lemire, S., Nawrocki, A., Kudirkiene, E., Møller-Jensen, J., Olsen, J. E., & Jelsbak, L. (2020). Polyamine depletion has global effects on stress and virulence gene expression and affects HilA translation in Salmonella enterica serovar typhimurium. Research in Microbiology, 171(3–4), 143–152. 10.1016/j.resmic.2019.12.001.

20. Han, J., David, D. E., Deck, J., Lynne, A. M., Kaldhone, P., Nayak, R., Stefanova, R., & Foley, S. L. (2011). Comparison of Salmonella enterica serovar Heidelberg isolates from human patients with those from animal and food sources. Journal of Clinical Microbiology, 49(3), 1130–1133. 10.1128/JCM.01931-10.

21. Hoffmann, M., Zhao, S., Luo, Y., Li, C., Folster, J. P., Whichard, J., Allard, M. W., Brown, E. W., & McDermott, P. F. (2012). Genome sequences of five salmonella enterica serovar Heidelberg isolates associated with a 2011 multistate outbreak in the United States. In Journal of Bacteriology (Vol. 194, Issue 12, pp. 3274–3275). 10.1128/JB.00419-12.

22. Huang, K., Wang, D., Frederiksen, R. F., Rensing, C., Olsen, J. E., & Fresno, A. H. (2018). Investigation of the role of genes encoding zinc exporters zntA, zitB, and fieF during Salmonella typhimurium infection. Frontiers in Microbiology, 8(JAN). 10.3389/fmicb.2017.02656.

23. Jelsbak, L., Thomsen, L. E., Wallrodt, I., Jensen, P. R., & Olsen, J. E. (2012). Polyamines are required for virulence in Salmonella enterica serovar Typhimurium. PLoS ONE, 7(4). 10.1371/journal.pone.0036149.

24. Lee, J., Monzingo, A. F., Keatinge-Clay, A. T., & Harshey, R. M. (2015). Structure of Salmonella FlhE, conserved member of a flagellar type iii secretion operon. Journal of Molecular Biology, 427(6), 1254–1262. 10.1016/j.jmb.2014.11.022.

25. Litvak, Y., Mon, K. K. Z., Nguyen, H., Chanthavixay, G., Liou, M., Velazquez, E. M., Kutter, L., Alcantara, M. A., Byndloss, M. X., Tiffany, C. R., Walker, G. T., Faber, F., Zhu, Y., Bronner, D. N., Byndloss, A. J., Tsolis, R. M., Zhou, H., & Bäumler, A. J. (2019a). Commensal Enterobacteriaceae Protect against Salmonella Colonization through Oxygen Competition. Cell Host and Microbe, 25(1), 128–139.e5. 10.1016/j.chom.2018.12.003.

26. Liu, W. T., Karavolos, M. H., Bulmer, D. M., Allaoui, A., Hormaeche, R. D. C. E., Lee, J. J., & Anjam Khan, C. M. (2007). Role of the universal stress protein UspA of Salmonella in growth arrest, stress, and virulence. Microbial Pathogenesis, 42(1), 2–10. 10.1016/j.micpath.2006.09.002.

27. Loui C, Chang AC, Lu S. (2009). Role of the ArcAB two-component system in the resistance of Escherichia coli to reactive oxygen stress. BMC Microbiol 9:183. 10.1186/1471-2180-9-183.

28. Mey, A. R., Gómez-Garzón, C., & Payne, S. M. (2021). Iron transport and metabolism in Escherichia, Shigella, and Salmonella. EcoSal Plus, 9(2), eESP-0034. 10.1128/ecosalplus.ESP-0034-2020

29. Misra, H. S., Maurya, G. K., Chaudhary, R., & Misra, C. S. (2018). Interdependence of bacterial cell division and genome segregation and its potential in drug development. In Microbiological Research (Vol. 208, pp. 12– 24). Elsevier GmbH. 10.1016/j.micres.2017.12.013.

30. Musa, L., Proietti, P. C., Marenzoni, M. L., Stefanetti, V., Kika, T. S., Blasi, F., Magistrali, C. F., Toppi, V., Ranucci, D., Branciari, R., & Franciosini, M. P. (2021). Susceptibility of commensal e. Coli isolated from conventional, antibiotic-free, and organic meat chickens on farms and at slaughter toward antimicrobials with public health relevance. Antibiotics, 10(11). 10.3390/antibiotics10111321.

31. Nair, A. V., Singh, A., Devasurmutt, Y., Rahman, S. A., Tatu, U. S., & Chakravortty, D. (2024). Spermidine constitutes a key determinant of motility and attachment of Salmonella Typhimurium through a novel regulatory mechanism. Microbiological Research, 281. 10.1016/j.micres.2024.127605.

32. Nair, A. V., Tatu, U. S., Devasurmutt, Y., Rahman, S. A., & Chakravortty, D. (2023). Spermidine facilitates the adhesion and subsequent invasion of Salmonella Typhimurium into epithelial cells via the regulation of surface adhesive structures and the SPI-1. bioRxiv, 2023-06. doi.org/10.1101/2023.06.03.543567.

33. Nair, D. V. T., Johnson, T. J., Noll, S. L., & Kollanoor Johny, A. (2021). Effect of supplementation of a dairy-originated probiotic bacterium, Propionibacterium freudenreichii subsp. freudenreichii, on the cecal microbiome of turkeys challenged with multidrug-resistant Salmonella Heidelberg. Poultry Science, 100(1), 283–295. 10.1016/j.psj.2020.09.091.

34. Nairz, M., Dichtl, S., Schroll, A., Haschka, D., Tymoszuk, P., Theurl, I., & Weiss, G. (2018). Iron and innate antimicrobial immunity-Depriving the pathogen, defending the host. In Journal of Trace Elements in Medicine and Biology (Vol. 48, pp. 118–133). Elsevier GmbH. doi.org/10.1016/j.jtemb.2018.03.007.

35. Nochino N, Toya Y, Shimizu H. 2020. Transcription factor ArcA is a flux sensor for the oxygen consumption rate in Escherichia coli. Biotechnol J 15:1900353. DOI:10.1002/biot.201900353

36. O’Connor, K., Fletcher, S. A., & Csonka, L. N. (2009). Increased expression of Mg2+ transport proteins enhances the survival of Salmonella enterica at high temperature. Proc. Natl. Acad. Sci. U.S.A. 106 (41) 17522–17527,

37. Pacello, F., Ceci, P., Ammendola, S., Pasquali, P., Chiancone, E., & Battistoni, A. (2008). Periplasmic Cu, Zn superoxide dismutase and cytoplasmic Dps concur in protecting Salmonella enterica serovar Typhimurium from extracellular reactive oxygen species. Biochimica et Biophysica Acta - General Subjects, 1780(2), 226–232. 10.1016/j.bbagen.2007.12.001.

38. Patterson, J. A., and K. M. Burkholder. “Application of prebiotics and probiotics in poultry production.” Poultry science 82, no. 4 (2003): 627–631. 10.1093/ps/82.4.627.

39. Pawitan, Y., Michiels, S., Koscielny, S., Gusnanto, A., & Ploner, A. (2005). False discovery rate, sensitivity and sample size for microarray studies. Bioinformatics, 21(13), 3017–3024. 10.1093/bioinformatics/bti448.

40. Rhen, M. (2019). Salmonella and Reactive Oxygen Species: A Love-Hate Relationship. In Journal of Innate Immunity (Vol. 11, Issue 3, pp. 216–226). S. Karger AG. doi.org/10.1159/000496370.

41. Russell, D. A., Dooley, J. S., & Haylock, R. W. (2004). The steady-state orgA specific mRNA levels in Salmonella enterica serovar Typhimurium are repressed by oxygen during logarithmic growth phase but not early-stationary phase. FEMS Microbiology Letters, 236(1), 65–72. 10.1016/j.femsle.2004.05.025.

42. Saldaña-Ahuactzi, Z., & Knodler, L. A. (2022). FoxR is an AraC-like transcriptional regulator of ferrioxamine uptake in Salmonella enterica. Molecular Microbiology, 118(4), 369–386. 10.1111/mmi.14970.

43. Samanovic, M. I., Ding, C., Thiele, D. J., & Darwin, K. H. (2012). Copper in microbial pathogenesis: Meddling with the metal. In Cell Host and Microbe (Vol. 11, Issue 2, pp. 106–115). 10.1016/j.chom.2012.01.009.

44. Schneider, B. L., Hernandez, V. J., & Reitzer, L. (2013). Putrescine catabolism is a metabolic response to several stresses in Escherichia coli. Molecular microbiology, 88(3), 537–550. 10.1111/mmi.12207

45. Schroll, C., Christensen, J. P., Christensen, H., Pors, S. E., Thorndahl, L., Jensen, P. R., Olsen, J. E., & Jelsbak, L. (2014). Polyamines are essential for virulence in Salmonella enterica serovar Gallinarum despite evolutionary decay of polyamine biosynthesis genes. Veterinary Microbiology, 170(1–2), 144–150. 10.1016/j.vetmic.2014.01.034.

46. Sun, C., Gao, X., Sun, M., Wang, Z., Wang, Y., Zhao, X., Jia, F., Zhang, T., Ge, C., Zhang, X., Zhang, M., Yang, G., Wang, J., Huang, H., Shi, C., Yang, W., Cao, X., Wang, N., Zeng, Y., Jiang, Y. (2022). Protective effects of E. coli Nissle 1917 on chickens infected with Salmonella pullorum. Microbial Pathogenesis, 172. 10.1016/j.micpath.2022.105768.

47. Taylor, A. L., Murphree, R., Ingram, L. A., Garman, K., Solomon, D., Coffey, E., Walker, D., Rogers, M., Marder, E., Bottomley, M., Woron, A., Thomas, L., Roberts, S., Hardin, H., Arjmandi, P., Green, A., Simmons, L., Cornell, A., & Dunn, J. (2015). Multidrug-resistant salmonella heidelberg associated with mechanically separated chicken at a correctional facility. Foodborne Pathogens and Disease, 12(12), 950–952. 10.1089/fpd.2015.2008.

48. Tripathi, A., Dewan, P. C., Barua, B., & Varadarajan, R. (2012). Additional role for the ccd operon of F-plasmid as a transmissible persistence factor. Proceedings of the National Academy of Sciences of the United States of America, 109(31), 12497–12502. 10.1073/pnas.1121217109.

49. Turnbough, C. L., & Switzer, R. L. (2008). Regulation of Pyrimidine Biosynthetic Gene Expression in Bacteria: Repression without Repressors. Microbiology and Molecular Biology Reviews, 72(2), 266–300. 10.1128/mmbr.00001-08.

50. Wang, Y., Yi, L., Zhang, J., Sun, L., Wen, W., Zhang, C., & Wang, S. (2018). Functional analysis of superoxide dismutase of Salmonella typhimurium in serum resistance and biofilm formation. Journal of Applied Microbiology, 125(5), 1526–1533. 10.1111/jam.14044.

51. Willis, L., & Huang, K. C. (2017). Sizing up the bacterial cell cycle. In Nature Reviews Microbiology (Vol. 15, Issue 10, pp. 606–620). Nature Publishing Group. 10.1038/nrmicro.2017.79.

52. Wu, S., Zhang, Q., Cong, G., Xiao, Y., Shen, Y., Zhang, S., Zhao, W., & Shi, S. (2023). Probiotic Escherichia coli Nissle 1917 protect chicks from damage caused by Salmonella enterica serovar Enteritidis colonization. Animal Nutrition, 14, 450–460. 10.1016/j.aninu.2023.06.001.

53. Zhao, X., & Drlica, K. (2014). Reactive oxygen species and the bacterial response to lethal stress. In Current Opinion in Microbiology (Vol. 21, pp. 1–6). Elsevier Ltd. 10.1016/j.mib.2014.06.008.

